# Cell-Free production of soybean leghemoglobins and non-symbiotic hemoglobin

**DOI:** 10.1101/2025.03.16.643390

**Authors:** Amanda P. Rocha, Mariele A. Palmeiras, Marco Antônio de Oliveira, Lilian H. Florentino, Thais R. Cataldi, Daniela M. de C. Bittencourt, Carlos A. Labate, Gracia M. S. Rosinha, Elibio L. Rech

**Author notes:** **Corresponding Author** Elibio Rech - Embrapa Genetic Resources and Biotechnology, National Institute of Science and Technology—Synthetic Biology, Brasília, DF, Brazil.

## Abstract

Hemoglobins are heme proteins and are present in some microorganisms, higher plants and mammals. In legume nodules there are two types: leghemoglobin (LegH) or symbiotic and non-symbiotic (nsHb). LegHs are present in high amounts at legumes roots and are responsible together with bacteroides for the nitrogen fixation process. Non-symbiotic hemoglobins Class 1 protein have very high affinity for O^2^ and are found in monocotyledons and legumes. LegH has aroused great interest in the vegetable meat industry due to its organoleptic and nutritional properties. Here, we demonstrated that soybean LegH A, C1, C2, C3 and nsHb are produced by *E. coli*-based cell-free protein synthesis (CFPS) and correctly synthesized in its amino acids sequence. In addition, it was also possible to reproduce some post-translational modifications confirmed by LC/MS analysis. All LegHs produced in this system showed peroxidase activity and heme binding correlated with its concentration in the assays. Furthermore, all proteins were readily digested by pepsin within 1 minute in analog digestion conditions. Therefore, LegHs and nsHb proteins were synthesized using cell-free systems (CFSs), maintaining their functionality and being digestible. These findings suggest that they could serve as viable alternative food additives for plant-based meat.

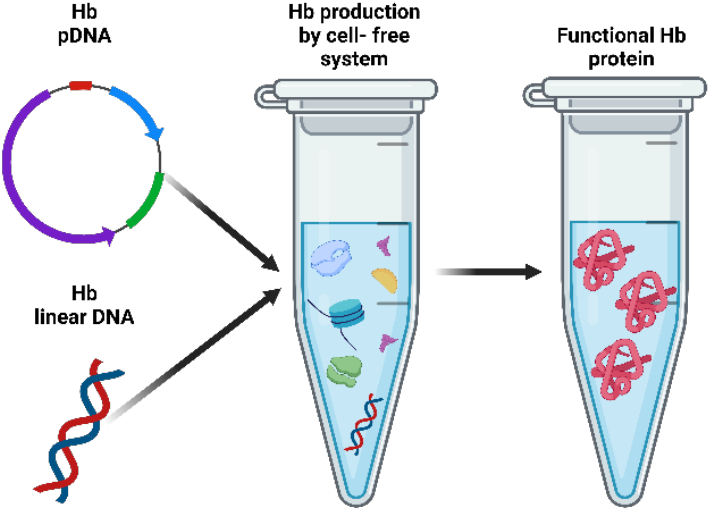

## 1. INTRODUCTION

Hemoglobins (Hbs) are heme proteins primarily responsible for the transport of O2. In plants, three types of hemoglobins have been identified: symbiotic hemoglobins, known as leghemoglobins (LegH), nonsymbiotic hemoglobins (nsHb), and truncated hemoglobins (tHbs).^1,2^ Leghemoglobin (LegH) is an essential protein for nitrogen fixation in the root nodules of legumes, promoting symbiosis between the host plant and bacteria of the genus Rhizobium^1^. LegHs were the first globin proteins discovered in plants and are found in high concentrations in the root nodules of soybean (*Glycine max*). In soybean there are five known genes that encode different LegH isoforms: LegH A, LegH Pseudogene, LegH C1, LegH C2, and LegH C3.^1,2^

In contrast to LegHs, which are specific to legume nodules, non-symbiotic hemoglobins are found in a wide variety of plant species and tissues. This includes soybeans, where nsHbs are expressed in various tissues such as the roots, leaves, and seeds, in addition to the symbiotic LegH found in nodules.^3^ While the exact roles of nsHbs are still being researched, studies suggest that they contribute to cellular energy production, especially under conditions of high energy demand or low oxygen availability.^4^ nsHbs are also expressed in the tissues of other plants, including the roots and seeds of rice, barley, and Arabidopsis, as well as in the leaves of alfalfa and the roots of cotton.^5^ This diverse expression pattern suggests that nsHbs may have different functions depending on their location and the specific needs of the plant.

Despite being discovered over 70 years ago^6^, LegH has recently garnered significant attention for its potential use as a food additive in plant-based meat products.^7^ This renewed interest stems from the growing demand for plant-based alternatives that closely mimic the sensory qualities of animal meat. Myoglobin, an iron-containing protein abundant in animal meat, plays a crucial role in the development of meat’s characteristic aroma, texture, and flavor during cooking. The heme group in myoglobin catalyzes certain reactions that transform amino acids, nucleotides, and sugars into complex flavor compounds. Recognizing this, the plant-based meat industry is exploring the use of heme-containing plant proteins, such as LegH, to replicate these desirable qualities in their products.^7^

In LegH, polypeptide chains bind to a heme B cofactor, which is identical to the heme found in animal meat and has been a part of the human diet for centuries.^8,9^ Importantly, the iron in LegH has the same bioavailability as the iron in bovine hemoglobin^10^ which is important considering that heme iron constitutes approximately 95% of the body’s iron store and is the primary source of iron for 67% of people in developed countries.^11^ Therefore, LegH presents a potentially valuable source of dietary iron while offering several advantages as a potential food additive. Its amino acid sequence does not share homology with known human allergens or toxins, and it is easily digested by pepsin under simulated gastric conditions.^12^

Impossible Foods produces soybean LegH C2 using the yeast *Pichia pastoris* as a host organism. This LegH serves as a key ingredient in their plant-based meat products, contributing to the characteristic flavor and aroma.^13^ The United States Food and Drug Administration (FDA) has authorized the use of this LegH as a color additive in plant-based burgers, with a maximum permitted level of 0.8%.^14,15^ The commercially produced soybean LegH C2, known as “LegH Prep”, has a purity of approximately 65%. The remaining 35% consists of other proteins derived from the *Pichia pastoris* host.^16^

Leghemoglobin has also been produced in *Escherichia coli*^17,18^, *Kluyveromyces marxianus*^19^ and *Corynebacterium glutamicum*.^*2*0^ However, in *E. coli*, the iron-containing heme groups in LegH can promote the formation of free radicals within the bacterial cells.^17^ This oxidative stress can damage cellular components and potentially hinder LegH production.

With respect to non-symbiotic hemoglobins, only a few have been expressed heterologously e.g. the type 1 and type 2 nsHbs from rice. A previous study aimed to elucidate the biological functions of non-symbiotic hemoglobins, which are thought play a significant role in the expression and utilization of oxygen in plants. The *in vivo* activities of rice nsHb-1 and nsHb-2 were investigated by analyzing their effects on the growth of *E. coli* TB1. The results revealed that growth inhibition was more pronounced when nsHb-2 was synthesized compared to nsHb-1, suggesting that these hemoglobins may have distinct roles *in vivo* within rice.^21^

Cell-free systems (CFSs) are rapidly emerging as a versatile platform for protein biosynthesis, offering a promising alternative to traditional cell-based methods. Unlike protein expression in living cells, which can be limited by cellular growth and complex regulatory networks, CFSs operate in a simplified environment.^22^ This approach eliminates the constraints imposed by cell membranes and complex cellular processes, allowing precise control over reaction conditions and the optimization of protein production.^22^ Moreover, these systems enable the production of proteins that may be toxic or difficult to express in living cells, such as membrane proteins and those with complex post-translational modifications.^23^

In this system, proteins are produced rapidly and efficiently using crude cellular extracts derived from prokaryotic or eukaryotic cells, which contain the essential native cellular transcriptional and translational machinery.^24^ The major components of cell-free protein synthesis reaction mixture are: template DNA encoding the target protein (circular or linear); a crude cellular extract; substrates for transcription and translation, including nucleotides and amino acids; and the components required for regeneration of ATP.^25^ Extract prepared from different organisms vary in terms of expression yield and the difficulty of synthesizing more complex proteins, for example eukaryotes which make post-translational changes, do not obtain high levels of protein. Therefore, expression systems that use prokaryotic organisms have a higher yield. However, these may not work in relation to some mammalian protein production, especially the ones requiring post-translational modification for proper function.^26^

Protein expression using CFSs has become very attractive compared to traditional methods, due to the ease with which proteins can be purified after expression, since the method has no cell wall, and the lysis stage is unnecessary. This makes the process much more agile and practical,^23^ while also offering the possibility of directing energy and metabolic expenditure only towards the production of the protein of interest, as energy expenditure towards the cell’s function and survival is not required.^27^ Finally, the cell-free expression method can produce proteins that would be lethal to the cell environment due to their toxicity, as they would inhibit transcription and translation, making the process of protein synthesis unfeasible.^26^

The increasing accessibility of CFSs is largely due to the availability of both commercial kits and home-made kits resources. Commercial kits, offered by various biotechnology companies which include cell extracts, enzymes, amino acids, and energy sources, alongside optimized protocols and DNA templates^28^, provide a standardized approach. On the other hand, the main advantage of producing home-made kits is the dramatic reduction of cost (up to 30x less^29^) while allowing greater flexibility and customization.^27^

In this work, we demonstrated for the first time that soybean leghemoglobins (LegH A, C1, C2, C3) and a non-symbiotic hemoglobin can be effectively produced using an *E. coli*-based cell-free protein synthesis system (CFPS). Additionally, these synthetically produced proteins showed peroxidase activity, possessed a heme group and were completely digested when exposed to simulated gastric fluid, thus increasing the number of heme proteins with potential use as additives in plant-based meat products.

## 2. RESULTS AND DISCUSSION

### 2.1. Cell-free synthesis of LegHs and nsHb proteins

Synthetic biology allows the optimization and simplification of biological processes, e.g., producing proteins in simple systems which mimic complex cells with only the essential machinery required for protein synthesis. This allows the substrates and energy to be directed only towards target protein production, in addition to permitting potentially toxic proteins to be produced, such as LegH. This protein possesses a heme group in its chemical structure containing iron atoms capable of producing free radicals, which cause cellular damage.^1^

Therefore, enabling its synthesis in synthetic biologic systems would be promising.

Four types of LegH were believed to derive from soybean nodules, LegH A, LegH C1, LegH C2 and LegH C3 and one nsHb^30^, however, a recent study has identified five LegHs and two nsHb in soy.^2^ Wiborg and colleagues analyzed the sequences and found that four of the LegH proteins have many conserved regions, explaining the similarity between them.^31^ The difference between the sequences of LegH A and LegH C3, the most distinct LegHs, is only 8%^31^, while around six amino acids differ in the sequences of the four LegHs analyzed (Figure S1). nsHb is found in soybean nodules and in other parts of the plant, such as embryos, leaves and roots^3,32^ and has a more distinct amino acid sequence compared to LegHs (Figure S1).

CFPS is a method based on only using the essential components needed to transcribe and translate DNA into protein in vitro, operating within a simple, open, and controlled system. This approach creates a versatile and easily manipulable system compared to living cells, enabling the addition of desired components while eliminating unwanted by-products that could inhibit protein synthesis. Furthermore, the transcription and translation capabilities of the CFS can accommodate various formats, including batch, continuous flow, and continuous exchange, thereby enhancing protein synthesis and increasing yields.^29^ Notably, CFS allow for the production of cytotoxic and transmembrane proteins in a controlled and optimized environment, which can be challenging to achieve in living cells.^33^ Among the various options, the *E. coli* machinery has demonstrated the highest efficiency in target protein synthesis.^29^

Four LegHs (A, C1, C2, C3) and nsHb DNA sequences from soybeans were synthesized into a pET28a vector, containing the promoter for T7 polymerase, RBS sequence and 6x Histidine tag at C-termini of each Hb (Figure S1). From these components, we have successfully produced all desired proteins using an E. coli based CFS prepared in our laboratory, operating for 16 hours at 28ºC. It is possible to see bands corresponding to LegHs (about 16 kDA with His-Tag tail) and nsHb (18kDA with His-Tag tail) by SDS-PAGE of raw CFS reaction extracts (Figure 1A). Electrophoresis of His-Tag purified proteins revealed two bands, one corresponding to our LegH (around 16-20 kDa) and one corresponding to the His-tagged T7 RNA polymerase included in the CFS reactions (∼ 100 kDa) (Figure 1A). The successful production of LegHs and nsHB by CFS and purification by affinity chromatography was also confirmed by immunoblotting analysis using antibodies against the 6xHis-Tag (Figure 1B). For purified fractions, we could identify the specific bands regarding LegH, also showing the efficiency of purification (Figure 1B and Figure 1C). We obtained an average of 0.5 mg/mL for both LegHs and nsHb in our optimized CFS.

**Figure 1.**
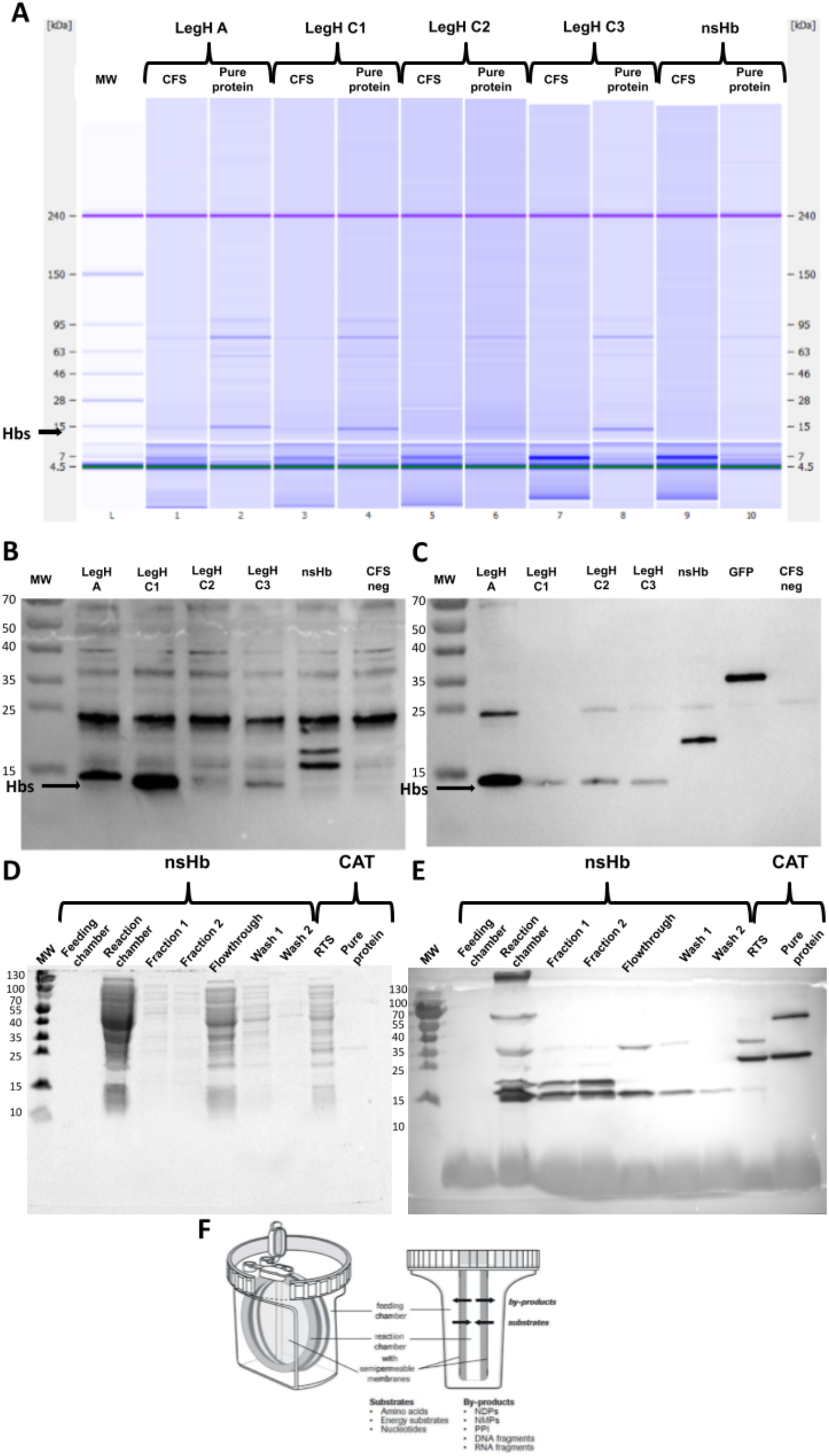
LegH A, C1, C2, C3 and nsHb production in a CFS. The reactions were performed as described in the methodology section using 20-50 nM of pDNA from LegH A, C1, C2, C3 or nsHb. Production of Hb proteins was confirmed in our CFS extract by protein electrophoresis of denatured proteins (A) and western blot using antibodies conjugated with HRP against 6xHis-Tag (B), giving protein bands at around 15 kDa. The 95 kDA band also seen corresponds to the T7 polymerase also recovered in the purification. The proteins produced were purified by affinity chromatography using nickel resin columns to obtain only His-Tag proteins for gel electrophoresis (A) and immunoblotting (C). nsHb was also produced by a medium scale commercial continuous-exchange CFS system (CECF) for 24 hours at 32ºC before being recovered and purified using affinity chromatography using a nickel resin. Samples were collected from the feeding and reaction chambers of CEFC device, as well as from elution fraction 1 and 2, the flowthrough and 2 steps of wash of chromatography column to analyze for the presence of nsHB. Recovered samples were analyzed by SDS-page and staining with Coomasie blue stain (D) and western blotting with anti-6xHis-Tag (E). Bands around 15 kDa representing nsHb and around 35 kDa representing CAT, used as the positive control supplied by the kit. The CECF device used and mechanism from Rabbit Biotechnology (#BR1400201) are pictured in (F). Device diagram reproduced from Rabbit Biotechnology brochure. **▾**

Since nsHb has often produced the most intense band in Western blot analyses, we wondered whether it can also be produced by medium-scale CFS. For this purpose, we used the commercial kit RTS 500 Proteomaster from Rabbit Biotecnhology (#BR1400201), which is based on a continuous-exchange cell-free (CECF) protein synthesis system (Figure 1F). This system consists of an inner chamber for protein prodution (reaction chamber), fed by an external chamber with an excess of substrates (feeding chamber) which also allows by-products to pass through the semi-permeable membrane, in order to not saturate the reaction chamber (Figure 1F).

nsHb was successfully produced in medium-scale CECF for the first time generating around 0.5 mg of target protein in 1 mL reaction with an *E. coli* extract. However, purification was unsatisfactory, because several bands were seen in SDS-page with Coomassie blue staining and in immunoblotting analysis, which also revealed the loss of proteins in wash steps (Figure 1D and 1E). The commercial CECF device from Rabbit Biotechnology used to produce is shown in Figure 1F. Therefore, optimization of chromatography to purify nsHB in medium-scale systems is required to improve production.

### 2.2 LC-MS analysis indicates that soybean LegHs and nsHB amino acids sequence was correctly synthetized in the cell-free system

Mass spectrometry analysis of the LegHs A, C1, C2, C3 and nsHb samples found 13, 20, 10, 14 and 7 peptides, covering 93%, 99%, 73%, 77% and 40% of the amino acid sequence respectively, with a false discovery rate (FDR) of 0,0% (Figure 2). Around 9, 9, 4, 2 and 6 unique peptides were found by LC-MS analysis in the LegH A, C1, C2, C3 and nsHb samples, respectively. It also identified the presence of post-translational modifications (PTM) in the Hbs, such as oxidation in LegH C1, C3 and nsHb (Figure 2). Our results show that CFS-produced LegHs and nsHb present correct amino acids sequence and some of their respective PTMs. However, we were unable to identify the PTM N-terminal acetylation^38^ in our synthesized LegHs, possibly due to peptide digestion prior to analysis. By analyzing our LC-MS results, we can identify the most common contaminating proteins present in our purified samples. The most common contaminant protein is the large ribosomal subunit protein uL3 (RL3), followed by the chaperone protein DnaK (DNAk) and T7 RNA polymerase (RPOL). Table 1 shows the nine most abundant proteins found in all or at least three or four experimental samples.

**Figure 2.**
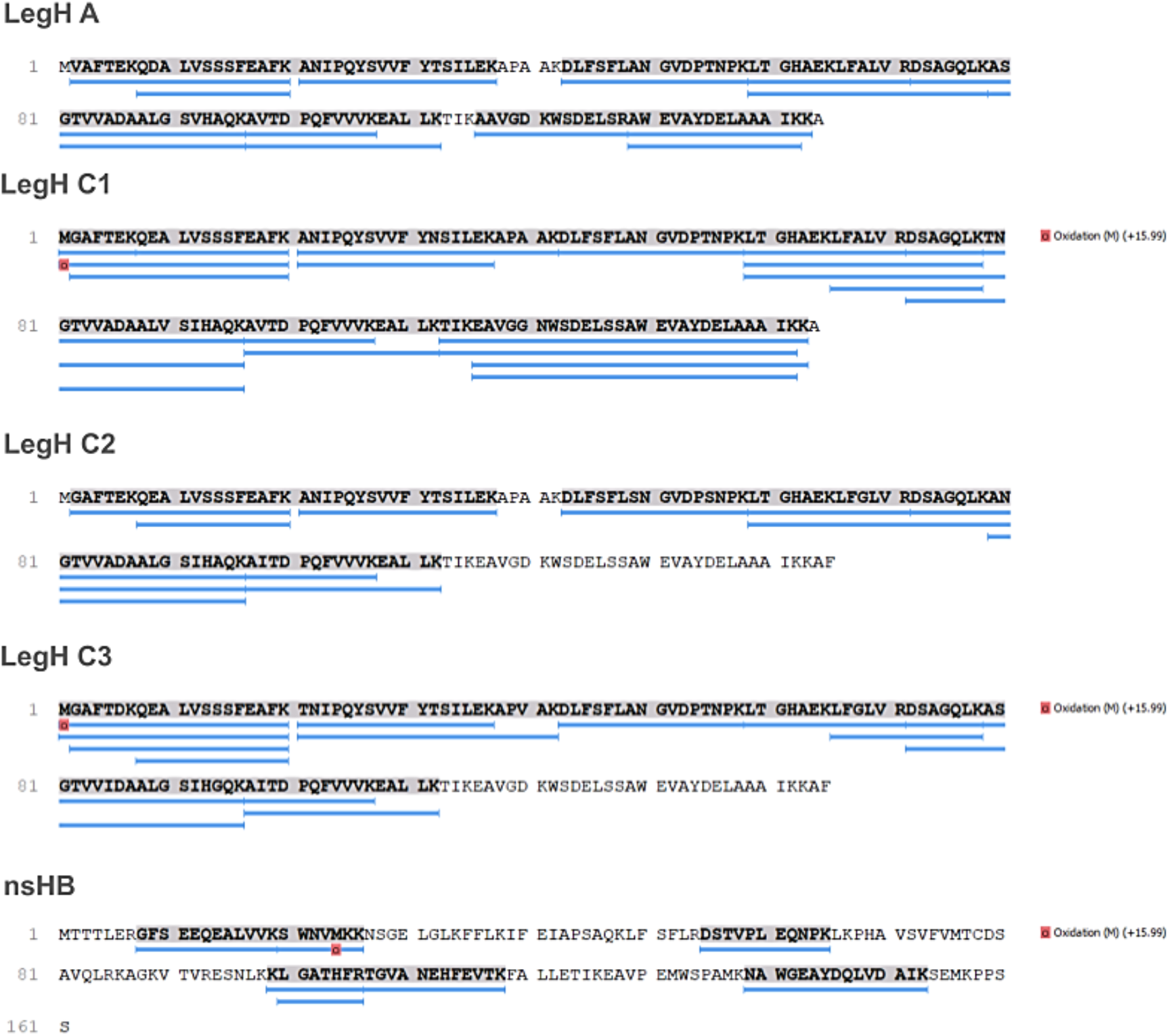
LegH A, C1, C2, C3, and nsHb were produced in a CFS with their correct amino acid sequences, as well as some post-translational modifications (PTMs). LegH A, C1, C2, C3 and nsHb were produced using 20-50 nM of pDNA per reaction, before being incubated at 28ºC, 800 rpm, for about 12-16h. Proteins were then purified, desalted, concentrated and digested by trypsin. The digested proteins were submitted to LC-MS and analyzed by Peaks BD software. Peptides detected from each sample were filtered using FDR up to 1% and aligned to the LegH A, C1, C2, C3 and nsHb amino acids sequences provided by the Uniprot database. The blue lines below each amino acid sequence indicate peptides found by LC-MS while the bold and highlighted letters represent which amino acids were identified. PTMs identified, such as oxidation, in LegHs and nsHb are shown as red squares under the respective amino acid.

### 2.3 LegHs and nsHB presented peroxidase activity

LegHs are known to present pseudo-peroxidase activity which may play a role in protecting against oxidant radicals in nodules forming compounds with hydrogen peroxide and reducing organic peroxides. ^19,39^ We verified that 40 µg/ml of synthesized LegH A, C1, C2, C3 and nsHb showed peroxidase activity in a dose-dependent fashion of around 7, 5, 2, 4 and 1 U/mL, respectively with LegH A showing the greatest activity. Therefore, 20-40 µg/mL of LegH and nsHb produced in CFS are sufficient to observe peroxidase activity. However, we were unable to detect peroxidase activity within 5-10 µg/mL of Hbs, suggesting an activity detection limit around these concentrations (Figure 3 and Figure S2).

**Figure 3.**
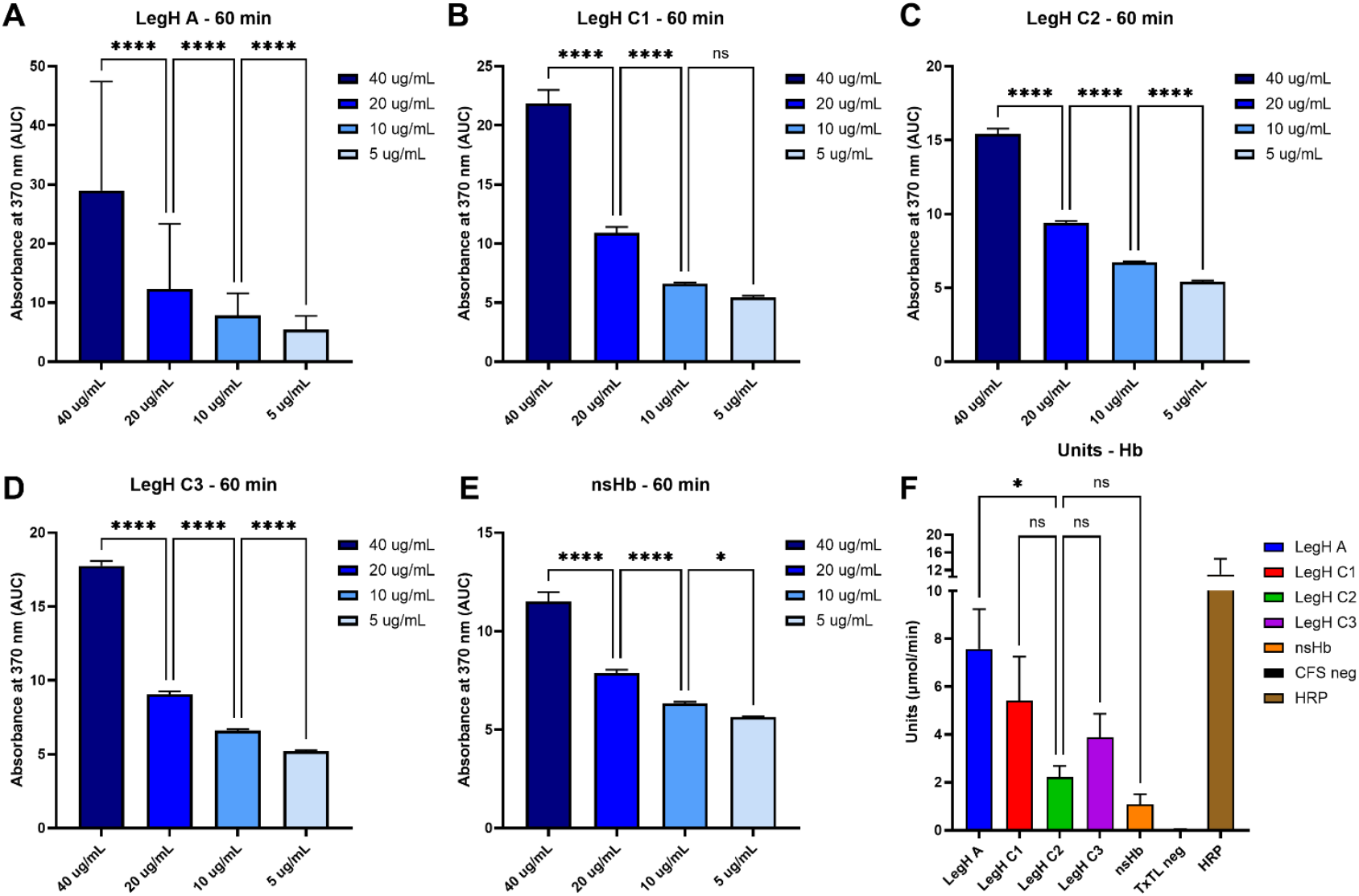
LegH A, C1, C2, C3, and nsHb produced from the CFS exhibited peroxidase activity. Purified Hbs were assayed for peroxidase activity over 1hr using TMB. The under-curve area was determined by the action of 40, 20, 10 and 5 µg/mL of LegH A (A), C1 (B), C2 (C), C3 (D) and nsHb (E). Units of enzymatic activity with 40 µg/mL of LegH A, C1, C2, C3 and nsHb (F). Values shown are means and standard error from four independent experiments performed in duplicate. P-values were calculated using one-way ANOVA with multiple comparisons corrected by Tukey’s test. Significance represented by * (p < 0,05); ** (p < 0,01); ***(p < 0,001); **** (< 0,0001). Ns: non-significative.

**Table 1.**
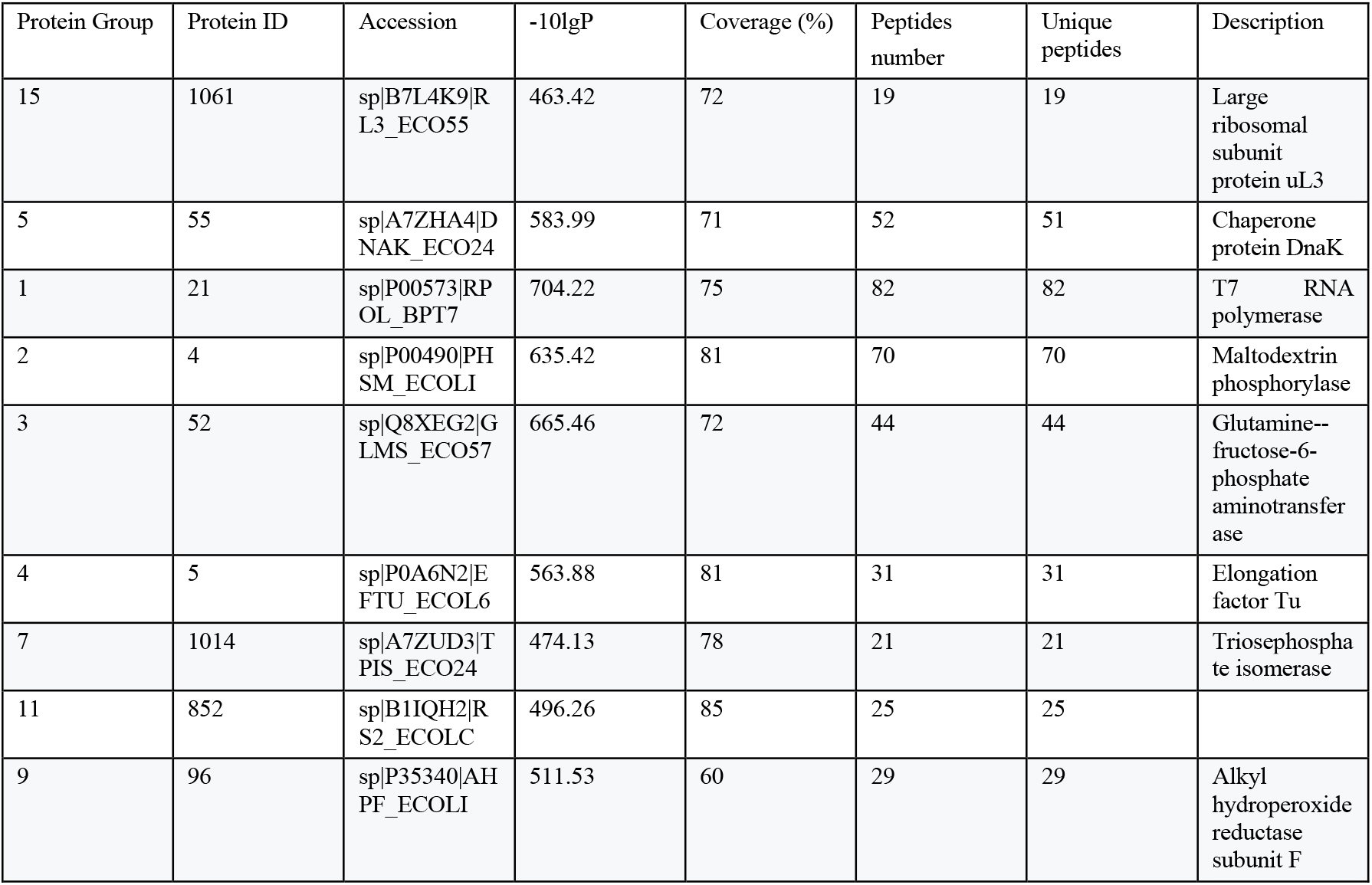
The most abundant proteins from *E. coli* in purified Hbs produced by the cell-free system found in LC-MS.

Soy LegH C2 produced in *P. pastoris* has been shown to present a peroxidase activity between 280 and 430 U/mg^40^ and Shao and colleagues attested that LegH C2 secreted by engineered *P. pastoris* presented a peroxidase activity of around 400 U/mg in a concentration of 250 mg/L, corroborating our results.^41^ LegH produced in *E. coli* has also shown peroxidase activity.^42^ Furthermore, comparing the LegHs produced in the cell-free system in our study, we observed that LegH A had almost four-fold more peroxidase activity compared to LegH C2, already used by the food industry to produce plant-based meat.

Other proteins have also been successfully produced by CFSs and are bioactive, such as insulin^34^, invasion plasmid antigens B (IpaB)^35^, serratiopeptidase^36^ and secretory leukocyte protease inhibitor (SLPI)^37^, suggesting this method provides native proteins production when the appropriate organism is used as extract source to provide some PTMs.

### 2.4 LegHs and nsHB produced from cell-free system contain a heme group

It is suggested that the heme group, similar to that found in myoglobin, becomes denatured and exposed when subjected to high temperatures during cooking. This denaturation promotes reactions that generate complex compounds responsible for the aroma, flavor, and texture of cooked meat. The presence of the heme group in soybeans and other legumes has sparked significant interest in the plant-based meat industry, driving efforts to create meat analogs that cater to vegetarian and vegan consumers.^7^

We confirmed that the LegHs and nsHb produced by CFS possessed a heme group. LegH A, C1, C2, C3 and nsHb showed a similar heme concentration in a dose-dependent fashion compared to Hbs concentration, with an average of 2 µM of heme with 60 µg of protein (Figure 4), corresponding to around 53% of heme-binding ratio as suggested by Shao and colleagues.^41^ Correlation coefficients using Pearson’s test between the heme and LegH ratio at different protein concentrations (30, 10 and 6 µg/mL) showed significant P values to LegH A, C1, C2, C3 and nsHb, of 0.034; < 0.001; 0.0024; 0.0058; 0.0365 and R squared of 0.9058; 0.9903; 0.9218; 0.8780 and 0.7050, respectively (Figure 4, inset). Except for nsHB, which showed a low R square, these data demonstrate that heme concentration decreased as LegH concentration also reduced.

**Figure 4.**
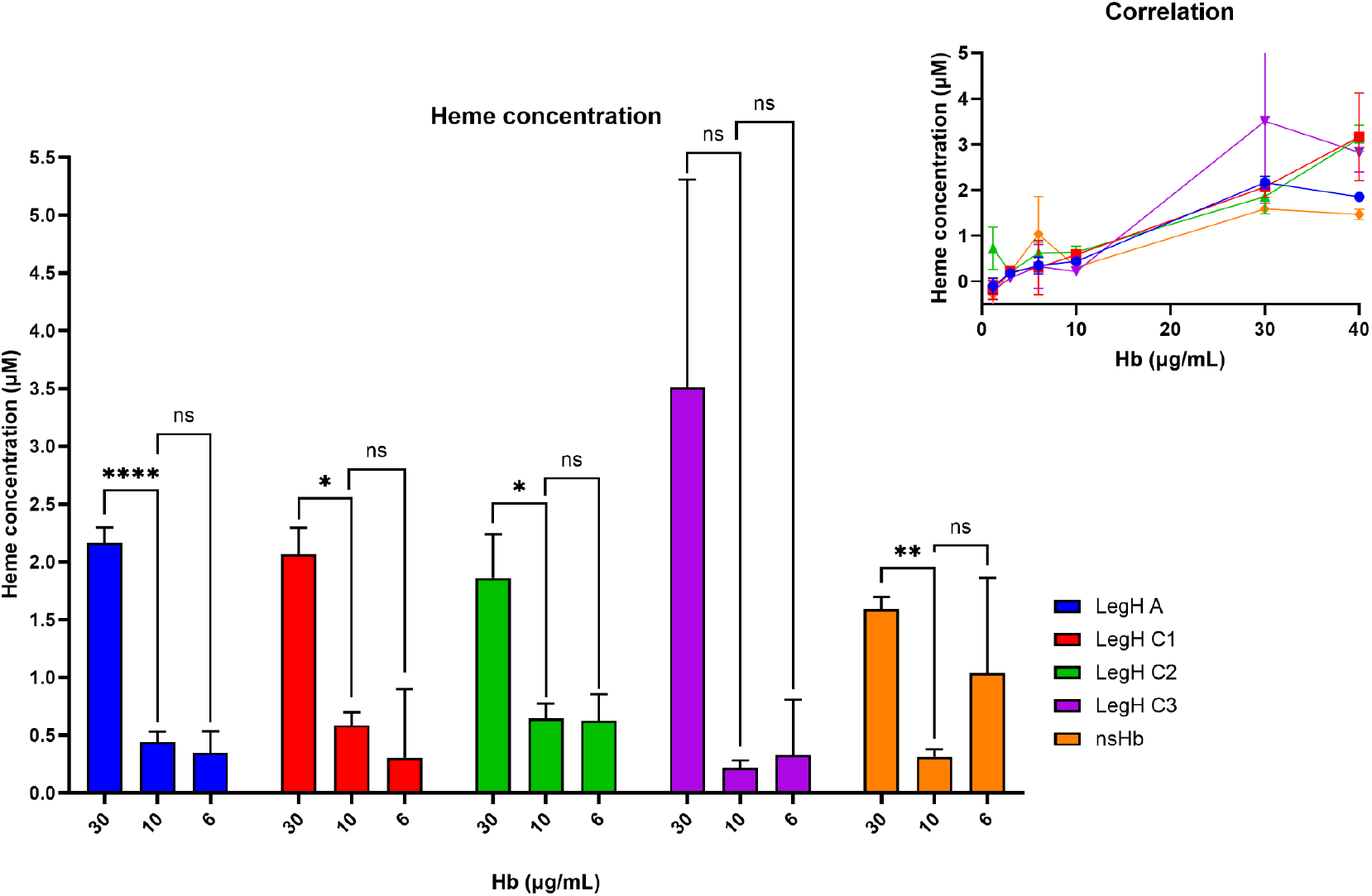
LegH A, C1, C2, C3 and nsHb produced from the CFS have a heme group in their structure. Purified Hbs were added to a heme reagent at a final concentration of 40, 30, 20 and 10 µg/mL of LegH A, C1, C2, C3 and nsHb. Heme concentration was measured from absorbance and was represented as the mean and standard error from three independent experiments performed in duplicate. Correlation graphic from heme concentration and hemoglobin proteins concentration (top right). P values calculated using one-way ANOVA with multiple comparisons corrected by Tukey’s test. Significance represented by * (p < 0.05); ** (p < 0.01); ***(p < 0.001); **** (< 0.0001). Ns: non-significative.

In this work we were also able to verify the presence of heme groups in other LegHs besides LegH C2 and nsHB. Until today, only LegHs purified directly from soybean root nodules^10^ and LegH C2 produced in *P. pastoris*^43,41^ were analyzed for the presence of the heme group. It is worth mentioning that heat process did not interfere in heme iron of LegHs^10^, suggesting that using these proteins as food additives may not be affected by cooking. It is possible that adjustment of the CFS allowing hemin supplementation could increase the heme concentration in the synthesized protein, in accordance with other studies which observed this increment at heme proteins.^44,45^

### 2.5 *In* vitro pepsin digestibility test

Protein digestion and degradation by pepsin have already been recommended to test allergenicity potential *in vitro* for extrapolation to human tolerance^46,47,48,49.^ For the analysis of the LegHs and nsHB produced in our CFS, they were incubated with pepsin in simulated gastric fluid, pH 2.0, simulating mammalian stomach conditions. All Hbs bands disappeared from our Western blots after 1 to 50 minutes of digestion with pepsin suggesting they may be digested under mammalian stomach conditions (Figure 5). This is corroborated with the previous findings, that attested both a LegH C2 preparation (mixture containing the soy LegH C2 isoform, residual *Pichia* proteins, and added food-grade stabilizers)^43^ and purified LegH C2^50^ was digested in 2 minutes by pepsin.

**Figure 5.**
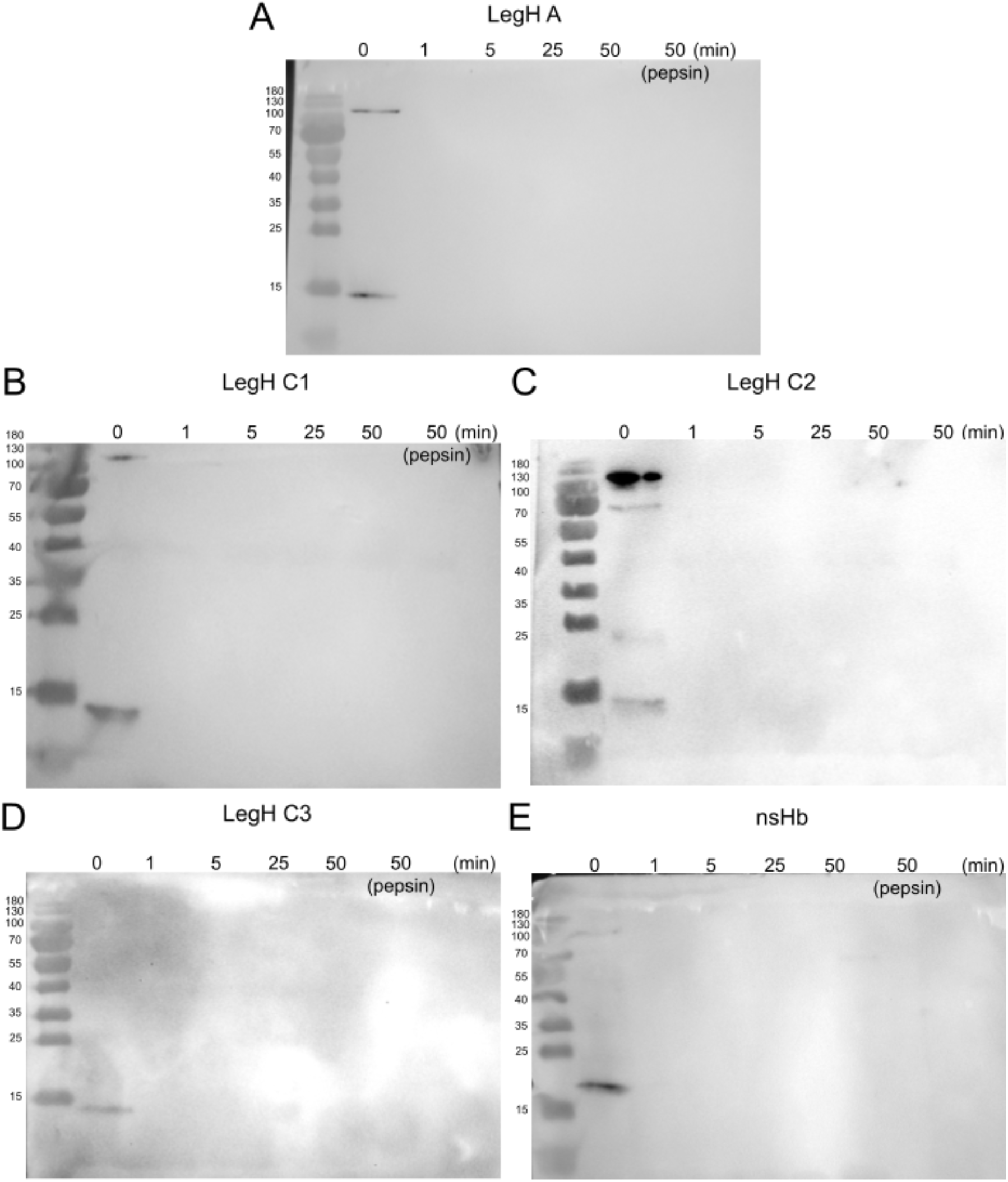
LegH A, C1, C2, C3 and nsHb produced in CFS systems are digested by pepsin. Hbs were incubated with pepsin (10 U/µg of LegH) in simulated gastric fluid (SGL) pH 2.0, at 37ºC, for 0, 1, 5, 25 and 50 min. After the defined times, SGL from each protein was recovered and neutralized with NaOHCO3 200 mM at pH 11 to inactivate pepsin action and stop digestion. These samples were denatured and submitted both to SDS-PAGE and immunoblotting analysis using anti-6xHis-Tag conjugated with HRP, then revealed with chemiluminescence (A-E).

## 4. CONCLUSION

Soybean LegH variants (C1, C2, C3, A) and nsHb can be efficiently produced using both commercial and laboratory-prepared cell-free systems (CFS), expanding the synthetic biology toolkit for generating these proteins essential to plant-based meat cultivation. These hemoglobins were successfully synthesized on small and medium scale, with the latter achieved through the CECF system. Proteomic analysis confirmed that LegHs and nsHb were produced with the correct amino acid sequences. Furthermore, LegH variants and nsHb synthesized via the cell-free system demonstrated pseudo-peroxidase activity and heme-binding capacity, highlighting their role in iron storage. All soybean hemoglobins studied were rapidly digested by pepsin within one minute, suggesting their potential as safe food additives for plant-based meats while minimizing the risk of intolerance. Therefore, we conclude that all tested LegHs and nsHb produced through the cell-free system retained the essential properties for their application as plant-based meat additives.

## 4. METHODS

### 4.1 Vector design and synthesis

The gene sequences of LegHs and nsHb were obtained from NCBI (LegH A Gene ID: 100527427, LegH C1 Gene ID: 100785236, LegH C2 Gene ID: 100527379, LegH C3 Gene ID: 100527391 and nsHb Gene ID: 102661758), codon optimized for *E. coli* and cloned into pET28a by EPOCH Life Science Inc (Missouri City, TX, EUA). Specifically, the genes were inserted between NcoI and XhoI restriction sites in the multiple cloning site of the plasmid pET28a (+). To facilitate protein capture and purification, LegHs and nsHb were expressed with a C-terminal His6-Tag.

Subsequently, the recombinant plasmids were transformed into DH5α *E. coli* strain, and a single colony was picked for expansion cultures. The plasmid DNA encoding LegH A, LegH C1, LegH C2, LegH C3 and nsHb were extracted with plasmid Maxi kit (Qiagen). Sequences and maps for all plasmids used are included as Supplementary file S1.

### 4.2 Cell extraction preparation

Cell-free extract and reagents used were prepared as previously described^51^, with necessary adaptations. Briefly, to obtain a cell-free extract, *E. coli* Rosetta 2 cells were initially cultured in 50 mL of 2x YPG medium supplemented with chloramphenicol at 37°C and 180 rpm for 16 hours. A 5 mL aliquot of this culture was then transferred to 750 mL of 2x YPG medium in a 2 L Erlenmeyer flask and incubated at 30°C and 180 rpm until they reached an optical density of 0.5 at 600 nm. The cells were harvested, washed, and lysed via sonication on ice (15% amplitude, 10 seconds on, 15 seconds off) with a total energy input of 2.7 kJ. The lysate was centrifuged at 15,000g for 30 minutes at 4°C. The supernatant containing the cell-free extract was collected and immediately frozen.

The amino acid mix was prepared using a 20 mM stock solution from each of the 20 standard amino acids: alanine, arginine, asparagine, aspartic acid, cysteine, glutamic acid, glutamine, glycine, histidine, isoleucine, leucine, lysine, methionine, phenylalanine, proline, serine, threonine, tryptophan, tyrosine, and valine. Each amino acid was dissolved in a 400 mM KOH solution adjusted to pH 6.6.

A 10X concentrated energy mix was prepared for the cell-free protein synthesis reaction. This solution contained 500 mM HEPES (pH 8.0), 15 mM ATP, 15 mM GTP, 9 mM CTP, 9 mM UTP, 0.68 mM folinic acid, 2 mg/mL E. coli tRNA mixture, 3.3 mM NAD, 2.6 mM coenzyme A, 15 mM spermidine, 40 mM sodium oxalate, 7.5 mM cAMP, and 300 mM 3-phosphoglyceric acid (3-PGA).

To acquiring T7 polymerase DNA, bacterial cells from *E. coli* strain BL21 previously transformed with the vector UMN 1396p (Pt7-911Q, generously provided by Kate Adamala – University of Minnesota) were cultured in 20 mL of LB medium with ampicillin at 37ºC, 180 rpm for 16h. Then, 5 mL of these cells were inoculated in 500 mL of LB medium containing ampicillin and incubated at 37ºC, 250 rpm until they reached an optical density of 0.5 at 600 nm when 1 mM IPTG was added into the culture. After 3 hours post induction, the culture was cooled and centrifuged for 10 min at 3700 rpm at 4ºC. Afterwards, the pellet was lysed with lysis buffer (50mM HEPES KOH pH 7.6, 1M NH4Cl, 10mM MgCl2), sonicated for 4 times (15% Amplitude (∼7W); cycles of 15s ON/15s OFF until 2kJ was reached) and centrifuged again for 45 min, 15.000g, 4ºC. The supernatant was recovered and purified in Ni-NTA agarose slurry in the chromatographic column. The purified T7 polymerase protein was dialyzed using Slyde-A-lyzer MWCO 30kDa cassettes and stored at -20ºC in 50% of glycerol until use. We confirmed purification of T7 polymerase by SDS-page and Western blotting, with a His-tag antibody confirming a protein of 106 kDa.

### 4.3 Cell-Free Protein Synthesis of LegHs and nsHb

Leghemoglobins and non-symbiotic hemoglobin were produced *in vitro* in a reaction composed of 12 mM M-glutamate, 140 mM K-glutamate, 1 mM DTT, energy mix diluted 1:10, 2 mM of each amino acid, 1 U/µL murine RNA inhibitor, 1.5 µM T7 RNA polymerase, cell free extract from *E. coli* Rosetta 2 diluted 1:3. The pDNA concentration generally ranged from 5 to 50 mM. The standard reactions were 50-100 µL with 16 hours incubation at 28ºC and stored at 4ºC short term or -20ºC for long term storage until used for downstream applications. GFP pDNA was used as positive reaction controls.

To produce nsHb on a medium scale, we used the CFS system from RTS 500 ProteoMaster E. coli HY (BiotechRabbit), which uses a continuous exchange cell-free (CECF) system, following the manufacturer instructions. 1 mL reactions were incubated at 32ºC, 800 rpm, for 20h. 20 µg of pDNA was used per reaction. Chloramphenicol acetyltransferase (CAT) vector was used as positive control.

For purification of LegHs and nsHB in small scale, we used His-Spin miniprep from Zymo Research (Cat # P2002) following the manufacturer instructions. For purification of LegH on a medium scale we used HisPur™ Ni-NTA Resin (Cat # 88221) according to the protocol from the producer. Briefly, incubated CFS reactions were mixed with Lysis buffer containing imidazole 10 mM, added to spin columns containing Ni-NTA resin and washed two times in the same buffer. Then, His-tag attached proteins were eluted by the addition of 300-500 µL of elution buffer with imidazole 300 mM. Purified proteins were dialyzed and concentrated using Amicon ultracel 3K. The proteins were centrifuged 3 times at 15.000 rpm for 20 minutes using 500 µL of PBS as exchange buffer.

### 4.4 Western blot analysis

To confirm the production of LegHs and nsHb, both whole CFS reactions and purified proteins were denatured in mPAGE 4X LDS buffer containing 25 mM β-mercaptoethanol at 70ºC for 5 min and submitted to polyacrylamide gel electrophoresis containing SDS (SDS-page), in a BIS-Tris gel with 15% of polyacrylamide. Electrophoresis was carried out for about 2 hours in a MOPs buffer. Then, proteins from the gel were transferred to a PVDF membrane using a semi-dry system for 30-40 minutes, applying 20 V. Following, the membranes were blocked with 5% BSA for 30 min at room temperature, incubated with anti-6x-His conjugated with either phosphatase alkaline (Cat # 46-0284) or HRP (Cat # A7058) at 4ºC, for 16h and then proteins were visualized using BCIP/NBT solution (Cat # 72091) 1:400 in Tris-HCL buffer pH 9.2 for 20-60 min or with Immobilon Forte Western HRP substrate (Cat #WBLUF0500) and revealed by chemiluminescence in iBright Image System (Invitrogen).

### 4.5 LC-MS and search parameters in public databases

Purified and desalted LegHs and nsHb were resuspended to a final concentration of 2µg/mL in 50mM ammonium bicarbonate with amicon ultra 0.5 mL. The proteins were denatured adding 0.025-0.1% RapiGest SF, vortexed and incubated at 80ºC for 15 min before adding 5 mM DTT and heated at 60ºC for 30 min. 15 mM iodoacetamide was then added before a second 30 min incubation before add-ing 1µg/µL of trypsin and the mixture was further incubated at 37ºC for 16h. Trifluoride acid was added to hydro-lyze RapiGest SF, before being centrifuged for 18,000g at 6ºC and the supernatant was recovered, concentrated and purified in a Reversed-Phase ZipTip C18, P10, (cat# ZTC18M096, Millipore). Samples were then resuspended in 0.1% formic acid and analyzed in a hybrid trapped ion mobility spectrometry-quadrupole time-of-flight mass spectrometer-timsTof Pro (Brunker Daltonics) assisted by chromatographic system nano Elute nanoflow (Bruker Daltonics) and an ion source CaptiveSpray. LC–MS was performed on an NanoElute (Bruker Daltonik) system coupled online to a hybrid TIMS-quadrupole TOF mass spectrometer^52,53^ (Bruker Daltonik tims TOF Pro, Germany) via a nano-electrospray ion source (Bruker Daltonik Captive Spray). For long gradients runs (2 hours total run), approximately 200 ng of peptides were separated on an Aurora column 25 cm × 75 µm ID, 1.9 um reversed-phase column (Ion Opticks) at a flow rate of 300 nL min-1 in an oven compartment heated to 50 °C. To analyze samples from whole-proteome digests, we used a gradient starting with a linear increase from 2% B to 17% B over 60 min, followed by further linear increases to 25% B in 30 min and to 37% B in 10min. Finally, to 95% B in 10 min which was held constant for 10 min. Column was equilibrated using 4 volumes of solvent A. The mass spectrometer was operated in data-dependent PASEF^54^ mode with 1 survey TIMS-MS and 10 PASEF MS/MS scans per acquisition cycle. We analyzed an ion mobility range from 1/K0 = 1.6 to 0.6 Vs cm-2 using equal ion accumulation and ramp time in the dual TIMS analyzer of 100 ms each. Suitable precursor ions for MS/MS analysis were isolated in a window of 2 Th for m/z < 700 and 3 Th for m/z > 700 by rap-idly switching the quadrupole position in sync with the elution of precursors from the TIMS device. The collision energy was lowered stepwise as a function of increasing ion mobility, starting from 20 eV for 1/K0 = 0.6 Vs cm-2 and 59 eV for 1/K0 = 1.6 Vs cm-2. We made use of the m/z and ion mobility information to exclude singly charged precursor ions with a polygon filter mask and further used ‘dynamic exclusion’ to avoid resequencing of precursors that reached a ‘target value’ of 20,000 a.u. The ion mobility dimension was calibrated linearly using three ions from the Agilent ESI LC/MS tuning mix (m/z, 1/K0: 622.0289, 0.9848 Vs cm-2; 922.0097, 1.1895 Vs cm-2; and 1221.9906, 1.3820 Vs cm-2).

Data processing, protein identification and relative quantification analyzes were performed using PEAKS studio Software, Version 10.6, Bioinformatics Solutions Inc., Waterloo, ON). Processing parameters included: cysteine carbamidomethyltion as fixed amino acid modification, whereas oxidation of methionine and acetylation of the N-terminal region were considered as variable modifications. Trypsin was used as a proteolytic enzyme, with a maximum of 2 possible cleavage errors. Minimum size for peptides were 7 amino acids. The ion mass deviation tolerance for peptides and fragments was set to 20 ppm and 0.05 Da, respectively.

A maximum false positive rate (FDR) of 1% was used to identify peptides and proteins, considering at least one unique peptide for protein identification as a criterion. All proteins were identified with a confidence level ≥ 95%, using the PEAKS Software algorithm and searching within the Uniprot database for Escherichia coli (taxon ID 562) and Glycine max (taxon ID 3847).

Proteomics results were filtered by Perseu software. Proteins in the matrix identified only by one modification site, as well as those identified by the reverse database and possible contaminants were excluded from subsequent analyses. Proteins were filtered to remain in the abundance matrix only those with values greater than zero in at least 50% of the samples from at least one of the groups. Afterwards, a script in the R programming language (https://www.R-project.org/) was used to refine the filter based on the percentage of protein presence in the groups and normalize the data by total ion count (TIC).

We also analyzed the contaminant proteins present at greater levels than our target proteins, those that were detected by LC-MS and presented the highest area peak on the mass spectrum. To this aim, we first classified the proteins that presented the highest peak area in decreasing order, thus the first proteins are the most present in our samples. The ten most abundant were selected from each sample, generating five tables with ten proteins, each one representing most presented in LegH A, C1, C2, C3 and nsHb. From these five tables, we observed which proteins were found in all or in three or four samples and then created a unique table with the most abundant proteins observed in LegH A, C1, C2, C3 and nsHb, overall. We filtered the proteins detected by those that presented the highest area of the peaks in decreasing order in each sample. Then we created a table with the contaminating proteins found more abundantly in all or 3 or 4 samples, respectively.

### 4.6 Assessment of LegH and Hb peroxidase activity

To verify whether LegH and nsHb presented pseudoperoxidase activity^55^ 10 µL of purified and dialyzed LegH and nsHb at 400, 200, 100 and 50 µg/mL were added to 100 µL of 3,3’,5,5’-Tetramethylbenzidine (TMB) liquid substrate system for ELISA (Cat # T0440 - Merck) in 96 clear flat-bottom well plates. The absorbance was then read at 370 nm in a spectrophotometer at 25º C every 1 minute for 60 min to confirm the peroxidase activity through the conversion of TMB into a blue product.

### 4.7 LegHs and nsHb heme binding

LegHs and nsHb heme binding was assessed by adding 50 µL of purified and dialyzed proteins, water (blank) or calibrator at 200 µL of heme reagent (Cat # MAK316 - Merck) in 96 clear flat-bottom well plates and the absorbance was measured at 400 nm at 25ºC. Heme concentration was calculated using the equation suggested by manufacturer instructions.

### 4.8 Pepsin digestibility assay

To assess the digestibility of LegHs and nsHb produced in our CFS system, we digested these proteins with pepsin (10 U/µg of LegH, Cat # P7012 - Merck) in a simulated gastric fluid (SGF) solution containing 0.084 N HCl and 35 mM NaCl, at pH 2.0 and 37ºC, for durations of 0, 1, 5, 25, and 50 minutes. After the defined times, 20 µL of SGL containing each protein were recovered and mixed with 7 µL of NaOHCO_3_ 200 mM at pH 11 to inactivate pepsin action and stop digestion. These samples were denatured and submitted to SDS-PAGE analysis as described above and digestion of LegHs and nsHb was evaluated.

### 4.9 Statistical analysis

Prism graphpad was used to plot data, create graphs and calculate statistical tests. Student T-Test and ANOVA two-tailed were used to calculate p value for quantitative data between two groups or three or more groups, respectively.

## AUTHOR INFORMATION

### Present Addresses

Amanda P. Rocha, Mariele A. Palmeiras, Marco A. de Oliveira, Daniela M. C. Bittencourt, Gracia M. S. Rosinha and Elibio Rech currently work at Laboratory of Synthetic Biology in Embrapa Genetic Resources and Biotechnology, Brasília, Brazil. Lilian H. Florentino currently works at Laboratory of Genetic Engineering Applied to Tropical Agriculture Biology in Embrapa Genetic Resources and Biotechnology, Brasília, Brazil.

Thais R. Cataldi and Carlos A. Labate currently work at Max Feffer Laboratory of Plant Genetics - EMU, Department of Genetics, Luiz de Queiroz College of Agriculture, University of São Paulo, Brazil.

### Author Contribution

A.P.R.: investigation, figure creation, data analysis, statistical analysis and manuscript writing. M.A.P.: investigation, manuscript writing. M.A.O.: implementation and optimization of the cell-free system in the Synthetic Biology Laboratory, writing - review & editing L.H.F.: investigation. D. M. C. B.: data analysis, writing - review & editing. G.M.S.R.: conceptualization, project administration, data analysis and writing - review & editing. E.R.: conceptualization, funding acquisition and writing - review & editing. T.R.C.: sample preparation and LC-MS analysis. C.A.L.: sample preparation, LC-MS analysis.

## ACKNOWLEDGMENT

This study was supported by Brazilian Agricultural Research Corporation (EMBRAPA, Brazil, and National Institute of Science and Technology in Synthetic Biology (INCT BioSyn - CNPq, Brazil, #465603/2014-9; FAP-DF, Brazil, #0193.001.262/2017 and #400145/2023).

## Supporting Information

Soybean Hbs sequences and peroxidase activity.pdf

